# Whole-brain deactivations precede uninduced mind-blanking reports

**DOI:** 10.1101/2023.04.14.536362

**Authors:** Paradeisios Alexandros Boulakis, Sepehr Mortaheb, Laurens van Calster, Steve Majerus, Athena Demertzi

**Author notes:** Corresponding author mail. Disclosure of competing interests: The authors declare no competing interests.

## Abstract

Mind-blanking (MB) is termed as the inability to report our immediate-past mental content. In contrast to mental states with reportable content, such as mind-wandering or sensory perceptions, the neural correlates of MB started getting elucidated only recently. A notable particularity that pertains to MB studies is the way MB is instructed for reporting, like by deliberately asking participants to “empty their minds”. Such instructions were shown to induce fMRI activations in frontal brain regions, typically associated with metacognition and self-evaluative processes, suggesting that MB may be a result of intentional mental content suppression. Here, we aim at examining this hypothesis by determining the neural correlates of MB without induction. Using fMRI combined with experience-sampling in 31 participants (22 female), univariate analysis of MB reports revealed deactivations in occipital, frontal, parietal, and thalamic areas, but no activations in prefrontal regions. These findings were confirmed using Bayesian region-of-interest analysis on areas previously shown to be implicated in induced MB, where we report evidence for frontal deactivations during MB reports compared to other mental states. Contrast analysis between reports of MB and content-oriented mental states also revealed deactivations in the left angular gyrus. We propose that these effects characterize a neuronal profile of MB, where key thalamocortical nodes are unable to communicate and formulate reportable content. Collectively, we show that study instructions for MB lead to differential neural activation. These results provide mechanistic insights linked to the phenomenology of MB and point to the possibility of MB being expressed in different forms.

## Significance Statement

This study explores how brain activity changes when individuals report unidentifiable thoughts, a phenomenon known as mind-blanking (MB). It aims to detect changes in brain activations and deactivations when MB is reported spontaneously, as opposed to the neural responses that have been previously reported when MB is induced. By means of brain imaging and experiencesampling, the study points to reduced brain activity in a wide number of regions, including those mesio-frontally which were previously detected as activated during induced MB. These results enhance our understanding of the complexity of spontaneous thinking and contribute to broader discussions on consciousness and reportable experience.

## Introduction

During spontaneous thinking, mental content appears continuous and seamless (Christoff et al., 2009). Probing people to report what they think yields various mental states with distinct contents and attitudes towards those contents, such as daydreaming, task engagement, and mind wandering (Van Calster et al., 2017; Smallwood et al., 2021). A critical component of these states is the presence of content. Recently, however, the study of unconstrained cognition has begun to focus on the experience of the inability to report on immediate mental content, termed mind blanking (MB; (Ward and Wegner, 2013)).

Recent research into the neural correlates of MB using fMRI experience-sampling (i.e., asking people at random times to report their immediate mental state, (Smallwood and Schooler, 2015; Weinstein, 2018) showed that spontaneous MB reports were close to a cerebral configuration characterized by a positive all-to-all connectivity profile (Mortaheb et al., 2022). Such a pattern of overall positive statistical dependencies implies that all cortical regions communicate in the same way when MB is reported. It is of interest that similar functional organization is observed in NREM sleep (El-Baba et al., 2019), suggesting that MB might be the result of overall low cortical arousal. Similar evidence was found on shorter timescales using EEG, where localized slow-wave activity was linked with MB reports, leading to the possibility of cerebral “local sleeps” during MB (Andrillon et al., 2019). Indeed, posterior electrode slow-wave activity during a go/no-go task was predictive of MB reports, in contrast to frontal electrode slow-waves which were linked to mind-wandering (Andrillon et al., 2021). Collectively, these studies propose that MB events are tied to neuronal profiles which do not permit efficient cortical communication, therefore hindering people from reporting clear mental content (Mortaheb et al., 2022).

A notable particularity of MB studies is the way MB is instructed for report. For example, Kawagoe et al. (2019) studied MB by asking people to actively “empty their minds” until they experience no thoughts, upon when they reported they had achieved this state. By analyzing the fMRI BOLD signal preceding these reports, the authors found deactivations in Broca’s area and the left hippocampus, and activations in the ventromedial prefrontal cortex (vmPFC)/ subgenual region of the anterior cingulate cortex (subACC). The authors interpreted these results as reduced inner speech, elicited by the attempt of participants to silence internally-generated thoughts. This possibility was considered before, primarily in the context of mind wandering. As our thoughts spontaneously transition across an internal-external milieu (Vanhaudenhuyse et al., 2011; Smallwood et al., 2012; Demertzi et al., 2013), the ACC serves executive functions, such as identifying attentional lapses from ongoing tasks (Christoff et al., 2009) or allowing thought transitions to be controlled (Crespo-Garćıa et al., 2022). In similar lines, self-induced MB also requires constant supervision of thoughts in the form of evaluating ongoing experience in order to promote thought-silencing, therefore recruiting regions such as the vmPFC/subACC, a central hub for mental state evaluative processes (Jenkins and Mitchell, 2011; Qin et al., 2020). For example, a hyper-experienced meditator showed decreases in fMRI connectivity between the posterior cingulate cortex and mesio-frontal regions when he was practicing content-free vs. content-related meditation (Winter et al., 2020). Taken together, the use of MB induction in neuroimaging studies might provide a biased picture about the underlying neural mechanisms of MB that incorporates task demands of thought monitoring.

In the present work, we test the hypothesis that uninduced MB reports are linked to frontal deactivations, inverting the pattern observed in self-induced MB. By means of fMRI and experience-sampling, we first performed a univariate analysis to test whether MB reports would indicate frontal deactivations in the periods preceding MB reports while remaining agnostic as to the contribution of the remaining cortex. Additionally, to supplement our hypothesis of frontal deactivations, we performed ROI analysis to examine the specificity of deactivations in the vmPFC-subACC and other previously identified MB-related clusters.

## Materials and Methods

### Experience-Sampling Dataset / Experimental Design

We used previously collected data (Van Calster et al., 2017) acquired during resting-state with eyes open in a 3T head-only scanner (Magnetom Allegra, Siemens Medical Solutions, Erlangen, Germany). At random intervals ranging from 30 to 60 seconds, participants were probed with an auditory cue to report via button press what was in their mind at the moment just preceding the cue. Each probe started with the appearance of an exclamation mark lasting for 1000 ms inviting the participants to review and characterize the cognitive event(s) they just experienced. After this period, participants were presented with 4 options, classifying their mental content as: a) Absence, defined as MB or empty state of mind, b) Perceptions, defined as thoughtfree attentiveness to stimuli via the senses, and c) Thoughts (Figure 1-A). In the case of a “Thought” report, participants were asked to report if the content was stimulus-dependent (sDEP; thoughts evoked from the immediate environment) or stimulus-independent (sIND; thoughts irrelevant from the immediate environment). Depending on the probes’ trigger times and participants’ reaction times, the duration of the recording session was variable (48-58 min). To minimize misclassification rates, participants had a training session outside of the scanner at least 24 hours before the actual session.

**Figure 1:**
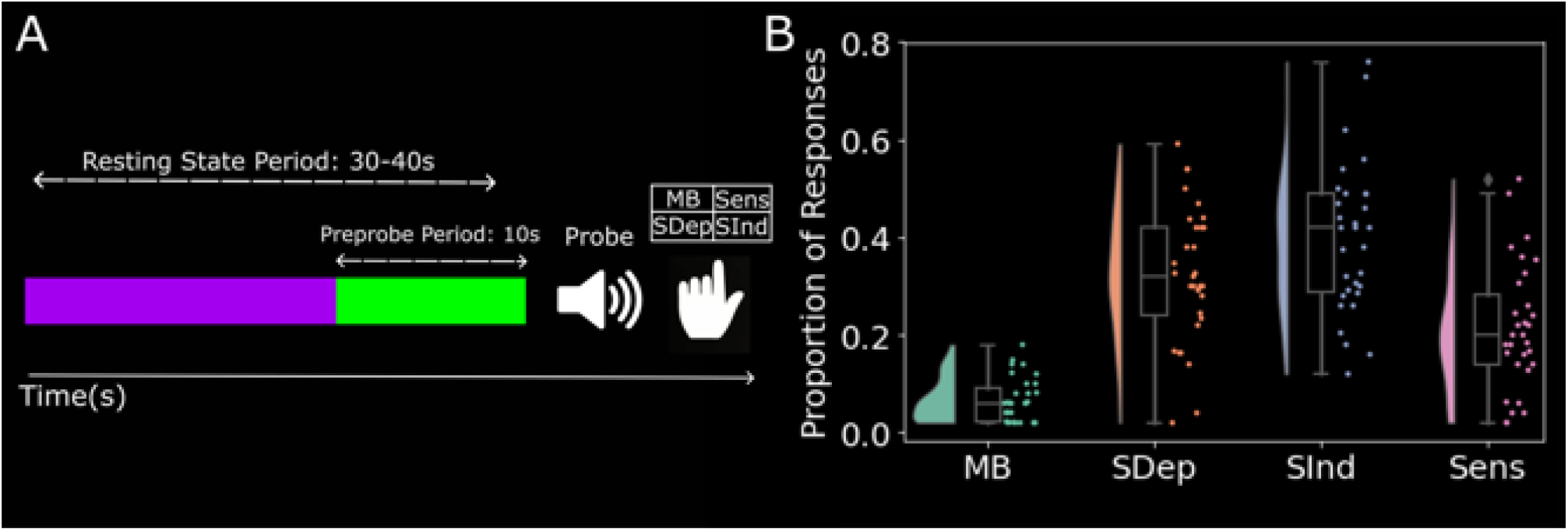
The Experience-sampling Paradigm. A. Single trial example: During experience-sampling participants are asked to restfully lay in the scanner with eyes open and let their mind wander without any further orientation as to the focus of their thoughts. At random intervals (30-60 sec), participants are probed with an auditory cue to report the content of their thoughts at the moment preceding the probe using button press. Four available report categories were presented as available: Mind-Blanking (MB), Sensations (Sens), stimulus-independent thoughts (sIND), and stimulus-dependent thoughts (sDEP). For subsequent analysis, only the final 10 seconds of the resting period (green segment) were used. B) Raincloud plots showing MB was reported at lower rates compared to mental states with content. Density kernels show how data are distributed and where peaks where aggregated. Boxplots show interquartiles ranges and medians. Pointplots show individual datapoints.

The dataset contains structural and functional MRI volumes for 36 healthy, right-handed participants (27 female, mean = 23, std = 3, range = [18,30]). Five participants were excluded as they did not report each mental state option at least once (total participants = 31, 22 female). Overall, participants reported MB 6% of total reports (SD: .04, Range: [1,9]) Sens 20% of trials (SD: .13, Range: [1,26]) SDep 32% of total reports (SD: .14, Range: [1,29]), and SInd 42% of total reports (SD: .15, Range: [6,28]). All participants gave their written informed consent to take part in the experiment. The study was approved by the ethics committee of the University Hospital of Lìege.

### fMRI acquisition parameters

fMRI data were acquired with standard transmit–receive quadrature head coil using a T2*weighted gradient-echo EPI sequence with the following parameters: repetition time (TR) = 2040 msec, echo time (TE) = 30 msec, field of view (FOV) = 192×192 mm2, 64×64 matrix, 34 axial slices with 3 mm thickness and 25% interslice gap to cover most of the brain. A high-resolution T1-weighted MP-RAGE image was acquired for anatomical reference (TR = 1960 msec, TE = 4.4 msec, inversion time = 1100 msec, FOV = 230×173 mm, matrix size = 256×192×176, voxel size = 0.9×0.9×0.9 mm). The participant’s head was restrained using a vacuum cushion to minimize head movement. Stimuli were displayed on a screen positioned at the rear of the scanner, which the participant could comfortably see using a head coil-mounted mirror.

### Statistical Analysis

#### Preprocessing

Structural and functional images were preprocessed using a locally developed pipeline written in the Nipype module (v1.8.2; https://nipype.readthedocs.io/) in Python (v3.8), combining functions from Statistical Parametric Mapping software (SPM12; https://www.fil.ion.ucl.ac.uk/spm/), the FMRIB Software Library v6.0 (FSL; https://fsl.fmrib.ox.ac.uk/fsl/fslwiki) and the Artifact Detections Tools (ART; https://www.nitrc.org/projects/artifact_detect). For each node of the pipeline, we have specified the respective module and function used. Structural images were skull stripped (fsl.Bet), bias-field corrected, and segmented into white matter, grey matter, and cerebrospinal fluid (spm.Segment). Finally, the restored, bias-corrected structural image was normalized into the standard stereotaxic Montreal Neurological Institute (MNI) space (spm.Normalize). The first four volumes (8.16s) of the functional data were removed to avoid T1 saturation effects (fsl.ExtractROI). The volumes were slice-scan time corrected to account for the accumulation of offset delays between the first slice and the remaining slices (fsl.SliceTimer). Then, the scans were realigned to the mean functional volume (spm.Realign) using a 2nd B-spline interpolation with leastsquares alignment. We used the realignment parameters to estimate motion outlier scans. An image was defined as an outlier or artifact image if the head displacement in the x, y, or z direction was greater than 3 mm from the previous frame, if the rotational displacement was greater than .05 rad from the previous frame, or if the global mean intensity in the image was greater than 3 SDs from the mean image intensity for the entire scans. The realignment parameters were also saved to a file so that these variables can be used as regressors when modeling subject-level BOLD activity. Then, the images were coregistered to the participant space using the bias-corrected structural image as the target and a normalized mutual information function (spm.Coregister) and then normalized to MNI space (spm.Normalize). Finally, the normalized images were smoothed using a Gaussian kernel of 8-mm full width at half-maximum. For comparability purposes, the preprocessing pipeline follows the approach as in previous works with this dataset(Van Calster et al., 2017) with MB analysis (Kawagoe et al., 2019).

#### Univariate whole-brain analysis

Data were analyzed using a univariate linear general linear model (GLM). The 4 responses of the participants (Absence, Perception, Stimulus Independent Thought and Stimulus Dependent Thought) were modeled and convolved with the canonical hemodynamic response function (HRF) as regressors of interest for each participant in the first-level analysis. Each response instance was modeled as an epoch starting 5 TRs before probe onset, following evidence from a “thinking aloud” paradigm that showed that mental states tend to fluctuate slowly, with one experience being reported every 10s (Van Calster et al., 2017). Each participant’s six motion parameters (three rigid body translations and three rotations from the realignment procedure) were included to regress out effects related to head movement-related variability. We used a high-pass filter cutoff of 1/128 Hz to remove the slow signal drifts with a longer period, and a first-order autoregressive model (AR (1)) was used for serial correlations with the classical restricted maximum likelihood (REML) parameter. Regionally specific condition effects were tested using linear contrasts for each key event relative to the baseline and each participant. Contrasts for “Perception” and “Thinking” regressors have been reported elsewhere (Van Calster et al., 2017). Therefore, here we tested for contrasts specific to MB. Given 4 regressors: [MB, Perception, sDEP, sIND], subject-level analysis yielded the following T contrasts of interest: a) Positive Effects of MB [1 0 0 0], b) Negative Effects of MB [-1 0 0 0], c) MB *>* Thinking [2 0 –1 –1], d) Thinking *>* MB [-2 0 1 1], e) MB *>* Perception [1 –1 0 0], f) Perception *>* MB [-1 1 0 0], g) Absence *>* Content [3 –1-1 –1], h) Content *>* Absence [-3 1 1 1]. The resulting contrast parameter estimates from the individual subject-level were entered into a random effects model for a second level analysis, using a one-sided, one-sample T-test. Regarding result reporting and visualization, we have opted for a “don’t hide/highlight” approach (Taylor et al., 2023). Effectively, we will be presenting all relevant maps at an uncorrected threshold of p_uncorrected_ *<* .001 Additionally, we will be annotating the contours of statistically significant clusters at a cluster-corrected threshold of pFDR*<*.05. Exploratory analysis will be conducted at clusters with voxel size *>*50. Interactive 3D surface projections of the contrasts presented in results are available on https://gitlab.uliege.be/Paradeisios.Boulakis/mb_activation/-/tree/main/plotting.

#### Region of interest (ROI) analysis

Based on the a-priori hypothesis about the role of the ACC in monitoring thought contents, we additionally performed a ROI analysis based on MNI coordinates reported in Kawagoe et al. (2019) for the ACC (MNI: 3,39,-5). To examine whether previous findings on the neuronal correlates of MB during active mental silencing can be extended to spontaneous blanking periods in ongoing mentation, we also included the left hippocampus (MNI: –27,-33,-3) and Broca’s area (MNI: –47,26,20).To extract single-participant beta parameters for each regressor of interest, 5-mm radius binary spheres were created for each ROI using the flsmaths function of the FSL software, which were then used to mask first-level subject-specific beta parameter maps, and extract the signal of interest. Localization of the ROIs was performed based on the MNI coordinates reported in Kawagoe et al. (2019) (Figure 4-A).

Given our hypothesis for the absence of frontal engagement in MB and the reduced statistical power of traditional frequentist approaches due to multiple comparisons, we opted for Bayesian Linear Modelling(McElreath, 2020), allowing us to make inferences on potential null results while not being overly conservative. For each ROI, we fit a linear model with beta values as a dependent variable, allowing the intercept to freely vary as a function of mental state.

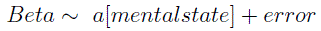

As prior for the intercept, we chose a normal distribution as it is the maximum entropy distribution (or “least surprising”) for any random variable with an unknown mean and unknown, finite variance. Effectively, a maximum entropy distribution is the most probable distribution for a random variable, given the potential constraints placed on its parameters. We chose to model the intercepts as *N* (0, 1), as we expected small effects (Figure 4-A). To examine the robustness of our choice of priors, we constructed two variants of normal distributions, one skeptical distribution that reduces effect sizes close to 0 by having high precision, marked as low variance *N* (0*, .*5), and one lax prior, marked by low precision, permitting extreme effects *N* (0, 3). Prior predicative simulation for the skeptical prior places the mass of effect sizes of each state within half a standard deviation from the mean. Likewise, the lax prior places the mass of effect sizes within 3 standard deviations. Additionally, we also fit a model using a uniform prior *U* (*−*2, 2), giving equal probability to effect sizes within two standard deviations from the mean. To further validate that our priors generated the desired ranges of parameters, we sampled from the prior distribution to perform a prior predictive visual check.

We estimated one posterior distribution for each one of the intercepts of the 4 mental states. Difference posterior was estimated by the pairwise subtraction of the mental state intercepts. Posterior distributions are summarized by their median, their standard deviation, and the 95% highest density intervals (HDI), representing the 95% probability that the true parameter lies within that range. To validate that the posterior accurately represented a generative model of the data, we also performed posterior predictive simulations, to examine if the ranges of our model can encompass the different beta values.

To fit the models, we used a Markov Chain Monte-Carlo No U-Turn Sampler (MCMCNUTS). MCMC is a class of algorithms for sampling from an unknown posterior distribution. The sampler uses a stochastic, random-walk procedure to draw samples from a random variable, and then approximates the desired distribution by integrating across the sum of the drawn samples (Harrison, 2010). The NUTS sampler is the mechanism of effective sample generation. As MCMC is sensitive to its tuning parameters, NUTS facilitates the sampling process by providing good candidate points in the distribution for the algorithm to sample (Hoffman and Gelman, 2011). To examine the convergence of the models, we sampled the posterior from 4 different chains, and both visually inspected the traceplot for points in the sampling procedure where the sampler stuck and accepted a model only if its scale reduction factor was at 1.00 (Figure 4-B). The stability of estimates was evaluated using an effect sample size (ESS) *>* 10000. We sampled 5000 samples from the posterior, with 2000 samples as burn-in.

Each model was compared to a null model:

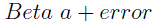

where the intercept does not differentiate between the mental states, effectively representing the mean of the mental states. Model fitting was performed using the PYMC3 (https://docs.pymc.io/en/v3/index.html) Python package Salvatier et al. (2016).

## Results

### fMRI univariate analysis reveals whole-brain deactivations

Initially, we focused on identifying regions associated with spontaneous MB occurrence during ongoing mentation. Overall, we found deactivations in the anterior cingulate cortex, the calcarine cortex, the bilateral thalami, the right anterior insula, the precentral gyrus, the left superior parietal lobule, the inferior frontal gyrus and the right operculum (Figure 2, Table 1). To validate these results, we examined different TRs around the probe period. Although an uncorrected voxel-level threshold of p = .01 recurrently showed deactivations in frontal, parietal and thalamical regions, cluster correction showed that only the thalamus was consistently deactivated across all time increments. Additionally, to control potential movement effects specific to conditions we estimated the overall framewise displacement of each subject at each timepoint (Power et al., 2012). Participants did not move significantly when considering displacement values per mental state category (MB: M = –.006, SD = .182, CI = [-.022, .009], SInd: M = –.003, SD = 0.143, CI = [-.008, .002], SDep: M = –.004, SD = 0.161,CI = [-.01, .003], Sens: M = –.006, SD = 0.254, CI = [-.018, .007]). Also, no significant difference was observed in terms of displacement values across mental states (F(1,4)=.146).

**Figure 2:**
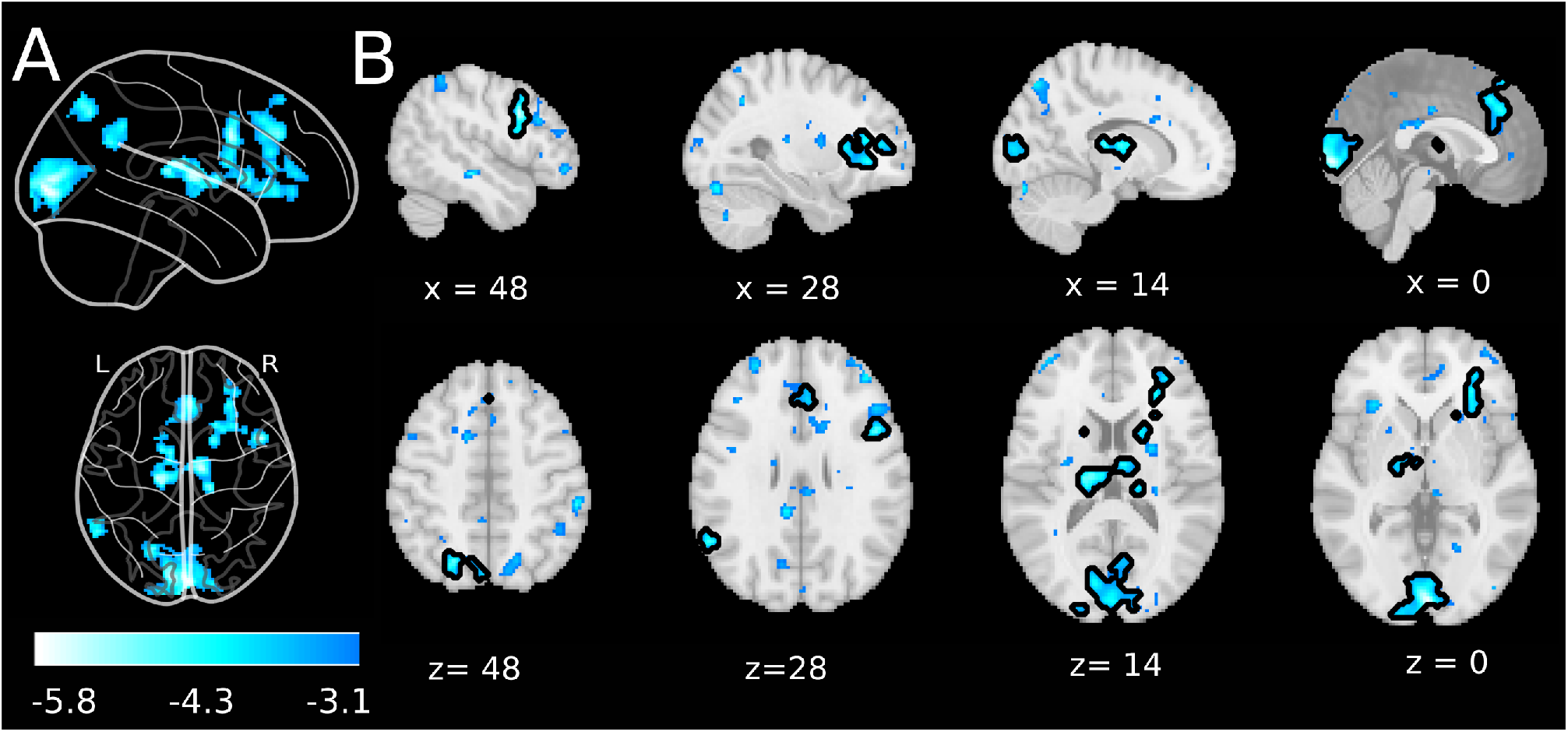
fMRI univariate analysis of MB reports reveals whole-brain deactivations. Statistically significant deactivations were observed in the anterior cingulate cortex, the calcarine cortex, the bilateral thalami, the right anterior insula, the precentral gyrus, the left superior parietal lobule, the inferior frontal gyrus and the right operculum. A) Glass brain projection (sagittal and axial views) at an at uncorrected voxel-level p_uncorrected_ *<* 001, and FDR-corrected at the cluster level p_FDR_*<*.05. Color-bar indicates T-statistic. B) Activation maps of negative MB effects projected on the MNI152 cortical template (sagittal and axial views). Maps are calculated on 10 seconds preceding MB reports. The deactivated map projection is performed at an uncorrected voxel-level p_uncorrected_ *<* .001 threshold level. Black contours signify the clusters that were significance at an FDR-corrected level of p_FDR_*<*.05.

**Table 1:** FMRI univariate analysis reveals deactivations during 5 TRs preceding MB reports.

| Region | No. of Voxels | Z peak | x | y | z |
| --- | --- | --- | --- | --- | --- |
| Right Calcarine Cortex | 1491 | 4.68 | 2 | -92 | 0 |
| Left Calcarine Cortex |  | 4.54 | -8 | -88 | 6 |
| Inferior Frontal Gyrus | 243 | 4.51 | 48 | 10 | 32 |
| Right Operculum |  | 3.76 | 48 | 10 | 22 |
| Right Thalamus | 617 | 4.49 | 12 | -12 | 8 |
| Left Thalamus |  | 4.49 | -18 | -18 | 16 |
| Superior Frontomedial Gyrus | 472 | 4.21 | 2 | 36 | 34 |
| Right Anterior Cingulate Cortex |  | 4.00 | 5 | 34 | 22 |
| Left Anterior Cingulate Cortex |  | 3.55 | -1 | 31 | 21 |
| Left Superior Parietal Lobule | 187 | 4.14 | -22 | -68 | 46 |
| Left Precuneus |  | 3.56 | -4 | -76 | 50 |
| Right Anterior Insula | 510 | 4.09 | 30 | 17 | 8 |
| Right Caudate |  | 4.09 | 18 | 8 | 16 |
| Left Supramarginal Gyrus | 164 | 3.99 | -58 | -56 | 28 |

*Note:* An uncorrected voxel-level threshold of $p = .01$ was set and FDR-corrected at the cluster level $p < .05$ . The X, Y, and Z coordinates refer to the AAL anatomical labeling map.

At the FDR cluster threshold (p*<*.05), the contrast between MB and the other mental states did not identify significant number of voxels. When the threshold was lowered to the exploratory level of whole-brain p*<*.001, voxels*>* 50, deactivations were observed in the angular gyrus (n voxels: 64, Z = 3.68, x = –60, y = –58, z = 32), a finding mainly driven by consistent deactivation of MB reports compared to stimulus dependent and stimulus independent thoughts (Figure 3). An examination of the individual regressor sign of activation (positive / negative) shows that MB tended to be significantly deactivated. On the other side, the other 3 mental states varied around 0, and as their confidence intervals included 0, we cannot clearly estimate the direction of their activation.

**Figure 3:**
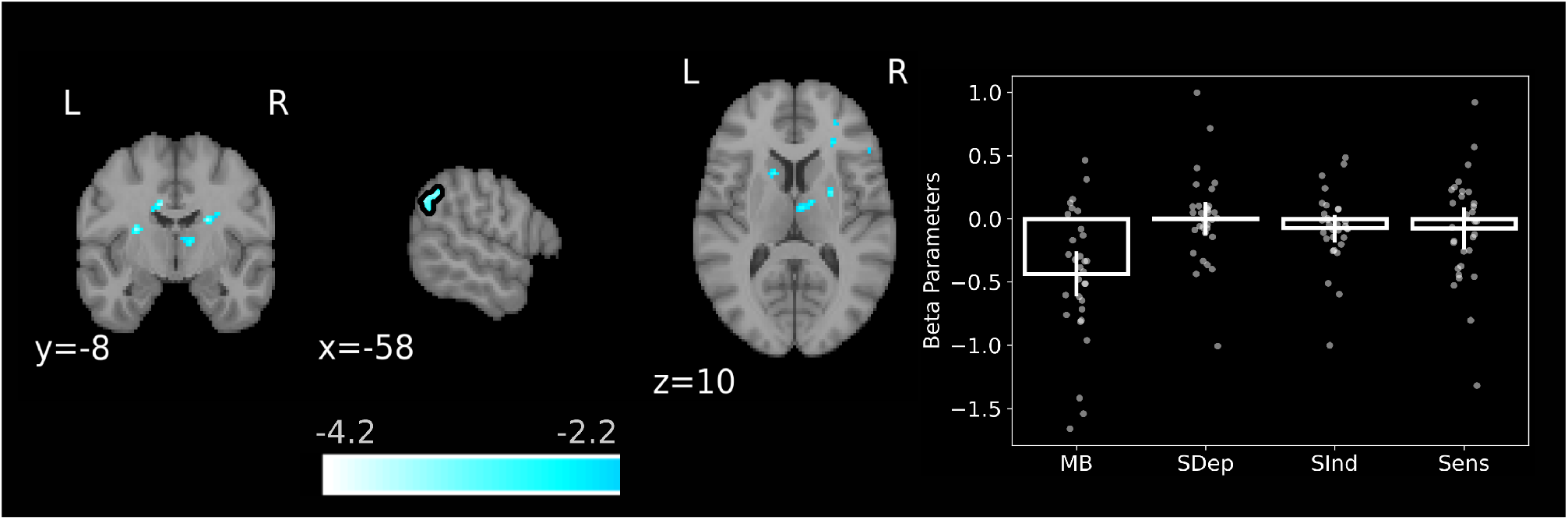
fMRI contrast analysis between content-oriented reports and MB at a lower exploratory threshold reveals deactivations in the left angular gyrus. (A) Activation map of presence vs absence of content contrast, projected on the MNI152 cortical template. The deactivated map projection is performed at an uncorrected voxel-level p_uncorrected_ *<* 001 threshold level. Black contours signify the clusters that were significance at an clusterextent theshold *<* 50 voxels. Color-bar indicates T-statistic. B) Boxplots representing the beta parameters of each mental state in left angular gyrus cluster. Error bars indicate 95% confidence intervals. Datapoints show single-subject parameter values.

### fMRI Bayesian ROI analysis provides evidence of deactivations in the vmPFC / subACC

Based on our a-priori assumptions about the role of vmPFC / subACC in thought monitoring, we examined the activation effects in the clusters reported in Kawagoe et. al, 2019, namely the vmPFC/subACC, Broca’s area and the left hippocampus. Extensive descriptive statistics of the posterior distributions for each ROI and mental state are presented in Table 2. Overall, the three ROIs’ MB intercepts did not include 0 in their 95% credibility intervals (vmPFC/subACC = Median: –0.242, SD: 0.119, HDI: [-0.471, –.01], Broca’s area = Median: –0.245, SD: .091, HDI: [-0.429, –.07], Left Hippocampus = Median: –0.113, SD: .056, HDI: [-0.219, –.001], (Figure 4-C), suggestive of functional deactivations in these clusters. To examine whether the clusters showed specificity in MB compared to the other mental states, pairwise comparisons between the MB beta parameters and the betas of each other mental state were calculated, as well as an overall MB vs. rest contrast. Pairwise comparison inference was performed by subtracting the MB posterior of each ROI from the posterior of the other mental states (Table 2). We found evidence only for the vmPFC/subACC cluster, namely MB reports were associated with reliably lower beta values compared to the other mental states (Median: –0.298, SD: 0.119, HDI: [-0.527, –.054]).Additionally, we found significant effects for the contrast MB-SInd (Median: –0.366, SD: 0.167, HDI: [-0.693, –.064])(Figure 4-E,G,I). Compared to the other mental states, MB was the only report category that was systematically deactivated, while the rest varied around 0. These results were consistent across the choices of different priors. No other ROI showed specificity for MB. To further validate whether the fitted models performed better against null models with only one intercept, for each fitted ROI we estimated the Watanabe-Akaike information criterion (WAIC) of the fitted and null model, as well as the expected log pointwise predictive density using leave-one-out cross-validation. Only for betas in the vmPFC/subACC did the model containing multiple intercepts perform better than the null model (Fitted_WAIC_ –129.687 *<* Null_WAIC_: –129.833, Fitted_ELPD_: –129.868 *<* Null_ELPD_: –129.908) (Table 3). The validity of the model fit, as well as the specificity of the vmPFC/ACC cluster in MB was replicated across all examined prior distributions for every model.

**Figure 4:**
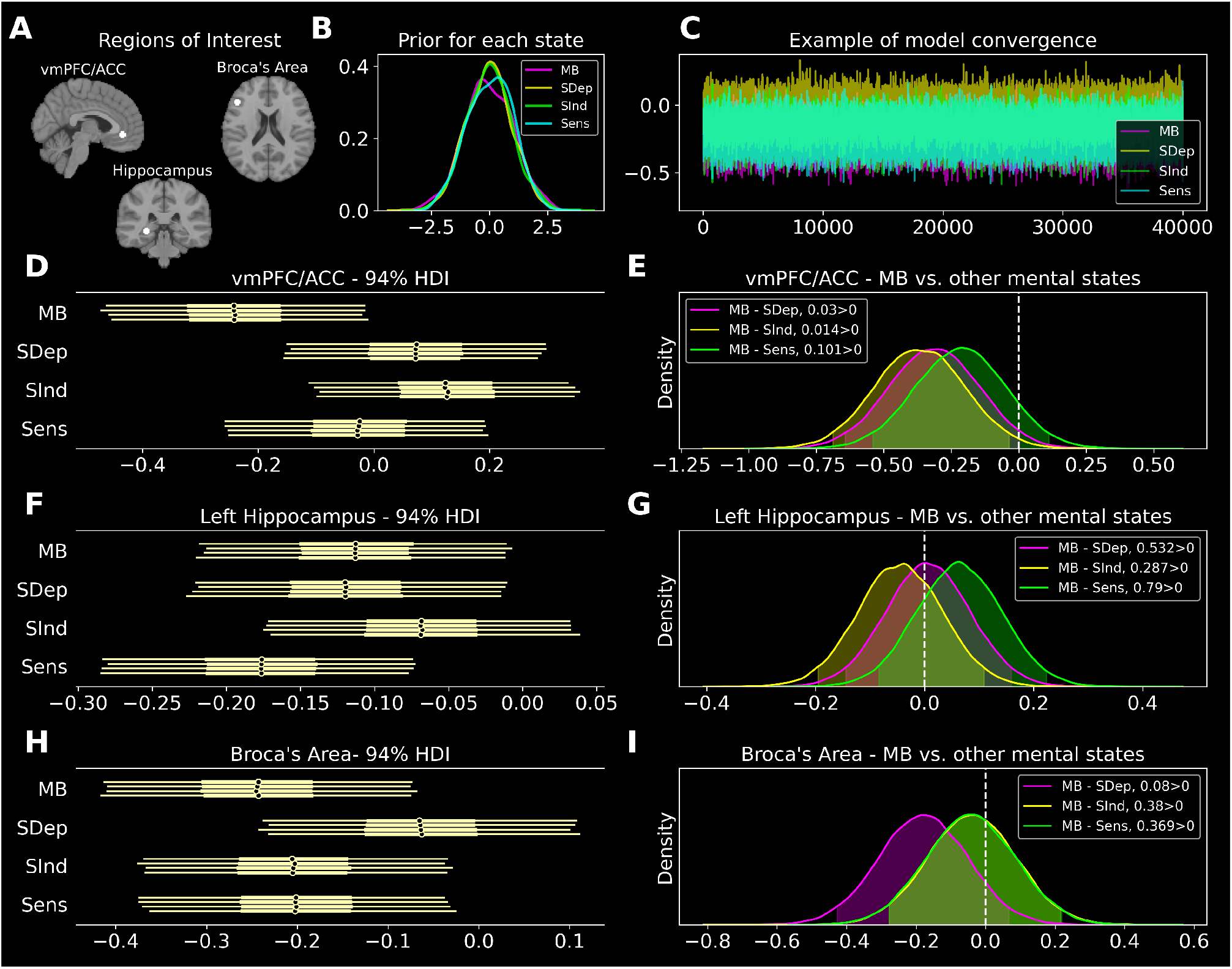
Bayesian analysis of the beta parameters in the vmPFC/ACC ROI reveals MB-related deactivations in this cluster. A) Regions of interest based on coordinates reported in Kawagoe et al. (2019) in the vmPFC/ACC, Broca’s Area, and the left Hippocampus. B) Null model prior expectations, modelling each prior as *N* (0, 1). C) Example traceplot of the model fit. Visual inspection of the random walk indicates that models converged, as the chains sampled the whole posterior space without autocorrelated data and sequential sampling of the same posterior space. D) Forest plot of each of the 4 sampled chains of the posterior distribution indicate that vmPFC/ACC contains significant evidence for MB deactivations, as the beta values did not contain 0 in the 94% HDI. Each line represents 94% highest density intervals (HDI). This was not the case for the rest of the mental states. E) Posterior differences between MB and the other mental states. We observed that the contrast MB-SInd did not contain 0 in the 94% HDI, providing evidence that frontal deactivations differentiated between MB and stimulus independent thoughts. F) Forest plot of each of the 4 sampled chains of the posterior distribution indicate that the left Hippocampus contains evidence for contribution only MB. This was also the case for SDep and Sens. G) Posterior differences between MB and the rest of the mental states at the left Hippocampus. We observed that no significant contrast indicating no specificity of the ROI in MB. H) Forest plot of each of the 4 sampled chains of the posterior distribution indicate that the Broca’s area contains evidence for MB contributions, as it does not contain 0 in the HDI. This was also the case for SInd and Sens. G) Posterior differences between MB and the rest of the mental states the Broca’s area. We observed that no significant contrast indicating no specificity of the ROI in MB.

**Table 2:**
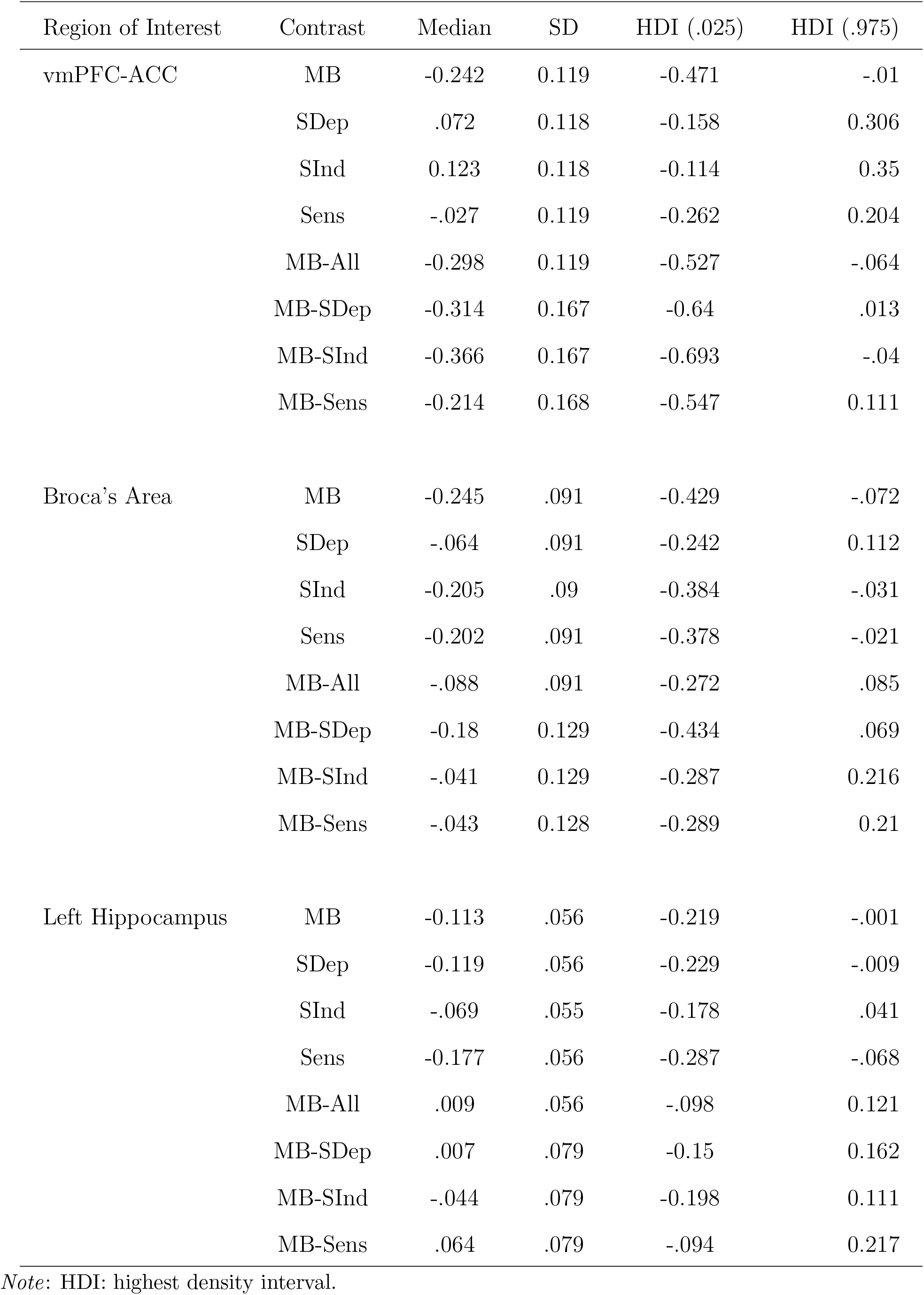
Descriptive statistics for the posterior distributions of the beta parameters for each ROI and mental state.

**Table 3:** Model comparison of fitted models.

| Region of Interest | Model | WAIC | ELPD |
| --- | --- | --- | --- |
| vmPFC-ACC | Fitted | -129.687 | -129.868 |
|  | Null | -129.833 | -129.908 |
| Broca's Area | Fitted | -93.934 | -93.972 |
|  | Null | -91.857 | -91.877 |
| Left Hippocampus | Fitted | -32.831 | -32.856 |
|  | Null | -30.82 | -30.847 |

*Note:* WAIC = Watanabe–Akaike information criterion, ELPD = Expected log pointwise predictive density.

## Discussion

We re-analyzed an fMRI experience-sampling dataset to study the neural correlates of mind blanking (MB) during unconstrained thinking and explore how instructions affect these correlates. Compared to mental states with reportable content, our findings indicate that spontaneous MB is linked to widespread deactivations in thalamo-cortical networks, which deviate from previous results.

### Widespread thalamo-cortical deactivations are linked to MB reports

We first show that whole-brain, thalamo-cortical deactivations precede MB reports. The fMRI univariate analysis, examining positive and negative effects of MB yielded deactivations in the anterior cingulate and calcarine cortex, the bilateral thalami, the right anterior insula, the precentral gyrus and the left parietal lobule. Such cortical deactivations have been previously associated with reduced neuronal resource allocation (Hester et al., 2004), task demands (Hairston et al., 2008) and impaired cognitive performance (Ji et al., 2010). Overall, we consider that the identified whole-brain deactivations might represent brief periods of neuronal disengagement, during which the brain cannot support attentional and mental-reporting processes.

This is further supported by the finding that two key subclusters were further deactivated: the primary visual cortex and multiple cortical nodes of the salience network (Seeley et al., 2007). In previous work, thoughts unrelated to the immediate environment correlated with the decoupling of sensory areas from regions contributing to stimulus salience (Mittner et al., 2016). Indeed, instructing participants to think of nothing results in decreased connectivity between the DMN and the sensory cortices, potentially reflecting this decoupling of the sensory system and a system of internal thoughts (Kawagoe et al., 2018). The whole-brain disengagement explanation is also supported by the deactivation of the thalamus, a recurrent node in saliency and engagement in mental state reportability (Kucyi et al., 2013). Thalamic activity covaries with executive control and attentional demands (Jansma et al., 2000; Antonucci et al., 2021). Potentially, the integrative nature of the thalamus (Hwang et al., 2017) is necessary to cast a mental spotlight and selectively allocate resources to bring a specific thought into conscious awareness. Overall, the rich profile of deactivations preceding MB reports highlights the important role of cortical nodes, traditionally associated with the salience of information.

On our quest to better understand the neuronal significance of such deactivations, we could resort to recent findings that analyzed the same dataset but examined functional connectivity. In that work, we show that MB reports are associated with a hyper-synchronized fMRI cortical connectivity profile, further characterized by high global signal amplitude, which we interpreted as neuronal down-states (Mortaheb et al., 2022). Although it would be tempting to hypothesize a similarly low neural activation mediating the identified deactivations, we recognize that a one-to-one comparison between the two analyses is difficult to make, as different aspects of the BOLD signal are examined. Indeed, while task-based BOLD activations can be considered as proxies of neuronal firing(Logothetis et al., 2001), changes in resting-state activity can result from complex interactions among neural, vascular, and metabolic factors (Liu, 2013). As a result, it is not clear whether there is a direct mapping between BOLD activations and functional connectivity analyses.

### MB-specific deactivations are linked to frontal and parietal regions

Moving to report-specific effects, by contrasting presence vs. absence of content we also found that MB is characterized by deactivations in the left angular gyrus. Supporting variant mnemonic (Ciaramelli et al., 2008), attentional (Cattaneo et al., 2009) and semantic processes (Kuhnke et al., 2022), the angular gyrus is recurrently present in content-oriented mental states. Indeed, angular activations have been correlated with both mind-wandering during ongoing mentation (Christoff et al., 2004; Maillet et al., 2019), and external orientation of thought during task engagement across demanding and non-demanding tasks (Turnbull et al., 2019). Therefore, the idea of generalized contributions of the angular gyrus to content-oriented mental states is further supported by our finding of inability to report mental content during deactivation of this region. Our results also are in line with previous electrophysiological results, where MB attentional lapses during task were predicted by posterior EEG slow-wave activity (Andrillon et al., 2021). The authors emphasized the role of parietal cortices in the emergence of conscious reports, where slow-wave activity might inhibit parietal-frontal communication and lead to the MB experience. We supplement this explanation by proving more granular structural information, introducing the angular gyrus as a an important parietal node.

By performing an ROI analysis to examine previously reported MB-specific cortical areas, we found MB deactivations in the ACC/vmPFC. In the context of thought-content, frontal activations were observed during mind-wandering with no meta-awareness compared to periods of mind-wandering with meta-awareness. The authors interpreted these larger activations as the ACC signaling a mismatch between expected thought stream and actual, wandering thoughts, eliciting a higher degree of surprise to the participant (Christoff et al., 2009). Additionally, vmPFC activation is correlated with episodic and social self-generated thought (Konu et al., 2020). However, given the multiple partitions of the ACC, treating it as a unimodal region that collectively contributes to one specific cognitive process might be misleading. In our study, the cluster originated close to the borders between ACC and vmPFC, denoting that the previous activation might include multiple processes (a detailed account can be found in https://neurosynth.org/locations/?x=4y=40z=-4). Indeed, the vmPFC-ACC cluster is systematically implicated in evaluative (D’Argembeau, 2013) and metacognitive processes (Vaccaro and Fleming, 2018), which are facilitatory to the internal stream of thought (Smallwood et al., 2012). Given the self-evaluative aspect of ACC-vmPFC, we here interpret these deactivations as failures to recurrently examine the content of a thought, which can be formulated as self-referential questions (“Am I thinking of anything?”) (D’Argembeau et al., 2007).

### MB as the mental state of “no thought”

A series of studies has explored ongoing thought using multi-dimensional experience sampling questionnaires, aiming to decompose it into a low-dimensional space where all content types can be represented (Mulholland et al., 2022; Konu et al., 2020, 2021). Interestingly, this approach has revealed an overlap in the low-dimensional space of ongoing thought-content between everyday life and in-lab task engagement, with consistent clusters related to social cognition, intrusive unpleasant thoughts, and task focus (Konu et al., 2021; Mulholland et al., 2022). In this space, where each dimension represents different content, we suggest that MB could represent the origin point, devoid of specific thought engagement, while moving away from this point would result in clearer content. Conversely, thoughts closer to the origin would exhibit less clearly reportable content. The activation patterns observed in the ventromedial prefrontal cortex (vmPFC) for thoughts along the social-episodic axis (Konu et al., 2020) and in the parietal lobule for thoughts along the task-focus axis (Turnbull et al., 2019) support this idea, as both these regions are deactivated during MB reports.

### Intentional and unintenional MB

So far, only one study has examined the fMRI neural correlates of MB from a univariate perspective (Kawagoe et al., 2019). In that protocol, participants were instructed to “think of nothing,” resulting in deactivations in Broca’s area and the left hippocampus, and activations in ACC. Similar frontal activations have been observed in clinical settings, where patients with depressive symptoms were guided to suppress their thoughts(Carew et al., 2015). By bridging the current literature together, we suggest that the discrepancy between uninduced and self-induced MB may reflect the existence of different forms of MB, similar to mind-wandering, for which intentional and unintentional forms have been proposed (Seli et al., 2016). Intentional MB may originate from top-down monitoring to exclude thoughts, such as meditation, while unintentional MB may arise from spontaneous lapses in frontal-parietal-sensory-thalamic systems that monitor the stream of consciousness and guide the ability to attribute semantic content to mental life. While this interpretation is still speculative and the clear presence of different MB forms cannot be extrapolated from our dataset, it paves a promising avenue for future research contrasting different forms of “thought absence”.

### Limitations and Conclusions

Several limitations pertain our study. The duration and sampling rate of mental states, including MB, in fMRI experience-sampling studies may lead to under-sampling of low-frequency and transient states (Mortaheb et al., 2022). Complementary methods, such as EEG, which allow for sub-second level estimation of brain dynamics, could provide valuable insights into momentary markers of MB. Additionally, the standard GLM-summary statistics approach may be suboptimal due to the fundamental unbalanced count of different mental states, resulting in reduced statistical power. In that sense, although the here identified effects remain safeguarded, we might nevertheless have missed others due to underpowered statistics. Finally, multivariable decoding approaches varying the duration of mental states could overcome the assumption of uniformity of mental state duration.

In conclusion, we investigated the neural correlates of uninduced MB during free-thinking conditions and found wide-spread thalamo-cortical deactivations, which may not allow the formulation of an efficient neural substrate to serve content reporting. We think that these results provide mechanistic insights on the phenomenology of MB and point to the possibility of MB being expressed in different forms. As MB holds experimental, philosophical, and potential clinical implications for understanding the thought-oriented and stimulus-driven mind, we believe future research would benefit by incorporating MB in the investigation of unconstrained thinking.

## Funding Sources

This article was supported by the Belgian Fund for Scientific Research (FRS-FNRS). It was also based upon work from COST Action CA18106, supported by COST (European Cooperation in Science and Technology).

## Code Accesibility

All codes to replicate the analysis is available on https://gitlab.uliege.be/Paradeisios. Boulakis/mb_activation (Boulakis, 2023). The code is based on existing Python libraries and custom functions. The provided repository contains all the necessary information to install an environment and reproduce the analysis on the experience-sampling dataset. We used an existing experience-sampling dataset, during which participants had the option to report the absence of thoughts (Van Calster et al., 2017). Previous research on this dataset, examining has replicated consistent fMRI findings in other mental states (MW: DMN and executive cortical areas). The raw data are also freely available in BIDS format from: https://openneuro.org/datasets/ds004134/versions/1.0.0. The unthresholded maps present in this paper can be found at https://identifiers.org/neurovault.collection:14761.

## Acknowledgments

We would like to thank Dr. Christophe Phillips, Dr. Federico Raimondo and Dr. Theodoros Karapanagiotidis for their meaningful discussions in the statistical analysis part of the manuscript

## Notes

https://openneuro.org/datasets/ds004134/versions/1.0.0

### Competing Interest Statement

The authors have declared no competing interest.

### Summary of Updates

The updated version of our manuscript contains conceptual clarifications of the term "Mind-Blanking", hence defined as "brief periods where people are unable to report their immediate-past mental state. Additionally, we have updated our preprocessing pipeline to be more methodologically sound. However, this does not affect our results.

